# Cognitive aging and categorical representations in visual working memory

**DOI:** 10.1101/2022.07.25.501371

**Authors:** Cherie Zhou, Monicque M. Lorist

## Abstract

A traditional view on cognitive aging is that visual working memory (VWM) capacity declines in older adults. Recent work has shown that visual information can be stored in VWM in different forms of representations. Specifically, VWM becomes more reliant on categorical representations (e.g., a prototypical red) as compared to continuous representations (e.g., a light reddish color) as memory load increases. Here, we replicated these findings and tested whether this holds for older adults. Participants memorized one to four colors; after a delay, an arrow pointed at the location of the color that needed to be reported. We used an extended mixture model (Zhou et al., 2022) to examine the extent to which memory responses were biased in the direction of the category prototypes. Our results showed that for both younger and older adults, VWM became more biased towards category prototypes with increasing memory load. Importantly, we found no difference in the extent to which VWM was biased towards category prototypes between younger and older adults. However, older adults showed an overall lower precision as compared to younger adults. Taken together, our results demonstrated that both younger and older adults became more reliant on categorical representations with increasing memory load; importantly, the extent to which categorical representations were involved in maintaining VWM information was insensitive to age.

---

Working memory provides a temporary storage system for information that is relevant to the task at hand (Baddeley, 2010; Oberauer, 2002). This information can be stored in the form of a precise, detailed representation, such as a specific shade of blue; or in the form of an abstract representation, such as blue as a general category. This distinction in the form in which visual working memory (VWM) can store information is commonly referred to as continuous and categorical representations, respectively.

Recent studies have shown that continuous representations are more fragile (Bae & Luck, 2019) and require more storage capacity as compared to categorical representations (Hardman et al., 2017; Zhou et al., 2022). This assumption is supported by the evidence that storing continuous representations appears to be more effortful than categorical representations, which limits the number of continuous representations that can be stored when memory load increases (Zhou et al., 2021). In a previous study, we asked participants to remember a varied number of colors (one to four); after a delay period of 100, 500, 1,000 or 2,000 ms, they had to report one of the memorized colors on a color circle (Zhou et al., 2022). Crucially, we measured a parameter that represents the extent to which VWM was biased categorically which we added to the original mixture model of WM (Zhang & Luck, 2008). We found that this categorical bias increased with memory load (see also Hardman et al., 2017; Panichello et al., 2019; Pratte et al., 2017), especially when the retention interval was longer than 1,000 ms. This increased reliance on categorical representations has also been shown when an ongoing VWM task is interrupted (Bae & Luck, 2019). Together, these findings suggest that VWM becomes more reliant on categorical representations when there is little capacity available for continuous representations.

Traditionally, older adults are considered to have severely limited VWM capacity as compared to younger adults, and therefore show lower memory precision (Brockmole & Logie, 2013; Hartshorne & Germine, 2015; Peich, Husain, & Bays, 2013). However, simply showing that older adults are worse in maintaining continuous representations says little about the complex VWM maintenance system (Perfect & Maylor, 2000). Recent work has shown that older adults may rely on different types of VWM representations in storing visual information in VWM than younger adults (Aziz, Good, Klein, & Eskes, 2021; Forsberg, Johnson, & Logie, 2020; Pilz, Äijälä, & Manassi, 2020). For example, in an orientation reproduction task, Pilz and colleagues (2020) asked the participants to remember a sine wave grating with varied orientations; after a masked delay, participants adjusted a bar to match the memorized orientation. Critically, the authors showed that older adults’ memory for oblique orientations (i.e. continuous representation) was substantially less accurate as compared to younger adults; however, their memory for cardinal orientations (i.e. categorical representation) was as accurate as younger adults. This suggests that although older adults have a declined capacity for storing continuous representations in VWM, they might still be able to keep visual information intact in VWM by relying on categorical representations.

So far, the distinctive role of continuous and categorical representation in aging has only been tested using indirect measures such as precision (Pilz, Äijälä, & Manassi, 2020) and probability of different types of representations (Forsberg, Johnson, & Logie, 2020). In the present study, we tested this question directly by measuring the extent to which VWM representations of color were biased categorically in both younger and older adults. We used a delayed estimation task (Ma et al., 2014), in which participants were instructed to memorize the exact shade of one to four colors; after a delay, an arrow was presented to indicate the location of one of the memory colors that they needed to report. Importantly, we implemented the adapted mixture model (Zhou et al., 2022) which characterizes the extent to which VWM relies on categorical representations. We fitted a crucial parameter from this model, the *categorical bias*, to our data; this parameter is coded in a way that positive values represent the response error in the direction of the prototypical color of the color category. A stronger categorical bias represents stronger reliance on categorical representations.

We predicted that if VWM relies more strongly on categorical representations with aging, then older adults will show a stronger bias towards category prototypes as compared to younger adults. Moreover, we predicted an effect of memory load on the extent to which VWM relies on categorical representations, such that VWM is more biased towards category prototypes with increasing memory load. We predicted no interaction between age and memory load.

## Method

### Preregistration

The detailed experimental designs and analysis plans were registered on the Open Science Framework (OSF) before we conducted the experiment: https://osf.io/hyxa3.

### Participants

We recruited 30 participants for both the younger (18-30 years old) and older (65-75 years old) group. This sample size is selected *a priori* based on our previous experiment which tested categorical bias using the same paradigm (Zhou et al., 2022). Participants were recruited from an online participant pool (Prolific, www.prolific.co), and completed the experiment online in exchange for a payment of £7.5 per hour. A total number of 37 younger and 46 older participants completed the experiment, from which 7 younger and 16 older participants were excluded based on the average rating criterion (see the criterion in *Data processing and exclusion criteria*).

All participants had normal or corrected-to-normal visual acuity and color vision, and indicated that they have no cognitive impairment or dementia. The study was approved by the local ethics review board of the University of Groningen (PSY-2122-S-0213). Participants provided informed consent before the start of the experiment. This experiment was conducted through an online server (JATOS; Lange, Kühn, & Filevich, 2015).

### Stimuli, Design and Procedure

As depicted in *Figure 1*, each trial started with a 500 ms memory display consisting of a number of colored circles (radius: 50 px). The number of circles varied from one to four depending on the Memory Load. These colored circles were evenly placed in a circular arrangement (radius: 250 px) around a fixation sign. The memory colors were randomly drawn from a HSV (hue-saturation-value) color circle with full value (i.e. brightness) and saturation for each hue. (Luminance ranged from 49 cd/m² to 90 cd/m² on a typical lab monitor. In the present study these values varied for each participant depending on their monitor.)

**Figure 1.**
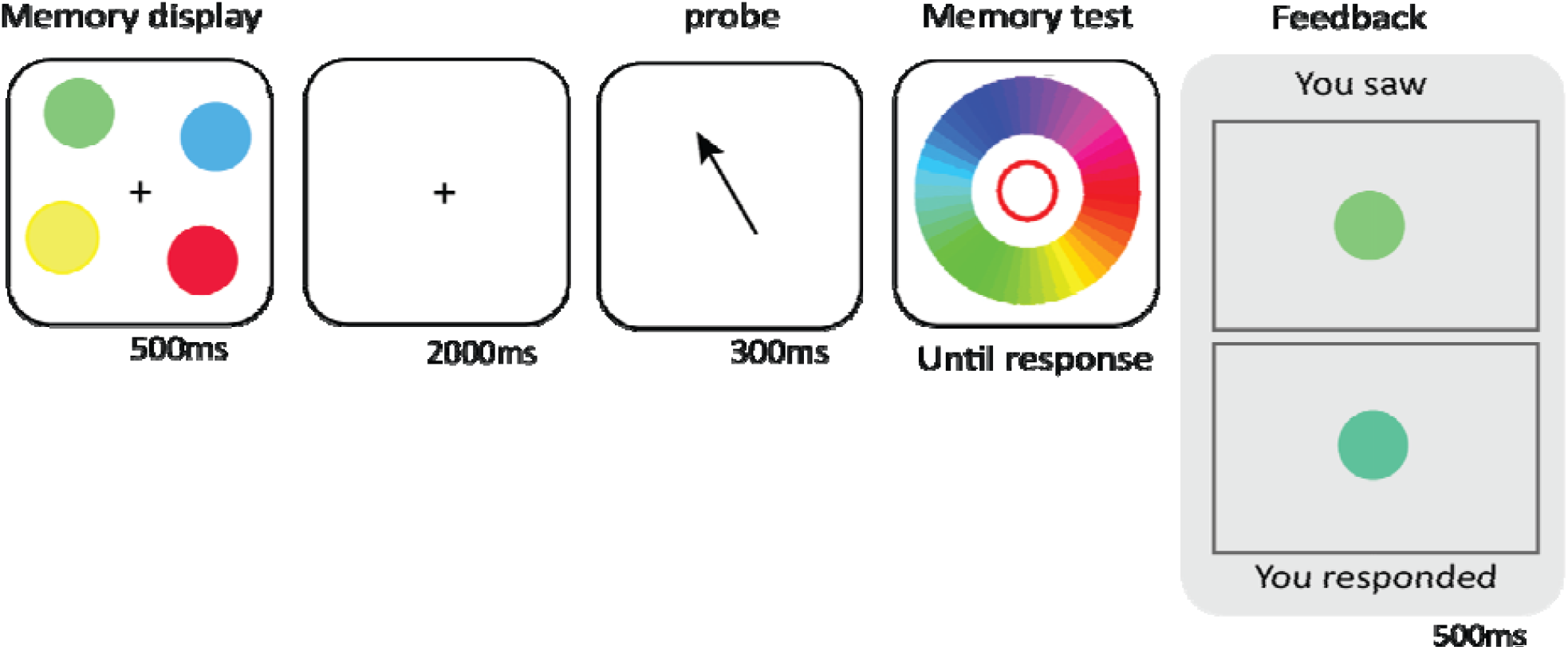
Sequence of events of a trial with a Memory Load of four.

Participants were instructed to memorize the exact shade of the colors of the circles. After a delay of 2,000 ms, a probe was shown for 300 ms, pointing at the location where one of the memorized colors was shown. Finally, participants selected the exact shade of the probed color on a color circle with no time limit. The color circle was randomly rotated on each trial. Each trial ended with visual feedback, showing both the actual color of the memory item and the color that participants reported. We used visual feedback to encourage non-verbal, visual processing and motivate participants throughout the experiment.

Following the main experiment, we asked participants to indicate the boundaries between color categories (e.g., the hue which they perceive as the boundary between green and blue) and the prototypical color for each color category (e.g., the hue which they perceive as the stereotypical color of green) on a color wheel. Each participant indicated each boundary and prototype twice. We used the following color categories: red, pink, purple, blue, green, yellow, and orange.

The four levels of the Memory Load conditions were mixed randomly within blocks without constraints. Participants completed 16 blocks of 25 trials each (i.e. 400 trials in total). Stimulus presentation and response collection was controlled with OpenSesame / OSWeb (version 3.3; Mathôt et al., 2012).

### Data processing and exclusion criteria

For each participant, we tested if the absolute error (i.e. the distance between the response hue and the target hue) was significantly below chance level (lower absolute error indicates better performance). To this end, we shuffled the response hue value of all trials for each participant, and then determined the “shuffled absolute error” based on the distance between the shuffled response hue value and the target hue value on each trial. Next, we tested if the response error of each participant was significantly lower than the shuffled response error with an independent sample t-test using an alpha level of .05.

For each age group, if any of the mean color-category boundaries or prototypical colors as rated by a participant deviated more than 2 SD^1^ from the grand mean ratings of the group, that participant wabe excluded. Excluded participants were replaced. Seven younger and sixteen older participants were excluded based on this criterion.

### Statistical Analysis

We calculated the response bias for each trial, which reflects the distance between the response hue and the target hue such that positive values reflect an error in the direction of the prototypical hue of the color category to which the target hue belongs. For example, if the target hue is an orange-like shade of red, then a positive response bias would indicate that the response hue was shifted in the direction of a prototypical shade of red. To take into account individual differences in color perception and the differences in color presentation on different monitors, we used the individually determined category boundaries and prototypes.

For each participant, we fitted an adapted mixture model (Biased Memory Model; Zhou et al., 2022) to the distribution of response bias for each memory load condition. This resulted in three parameters: 1) Bias ([-180, 180]): a parameter that reflects response bias such that positive values indicate a shift in the direction of prototypical colors; 2) Guess Rate ([0, 1]): a parameter that reflects the proportion of random responses; 3) Precision (or kappa; κ, [0, 10,000]): a parameter that reflects the standard deviation of the normal (von Mises) distribution, with a higher κ reflecting higher precision. We conducted three separate repeated measure analyses of variance (RM-ANOVA) with Age as a between-subject variable, Memory Load as a within-subject variable, and each of the three model parameters as dependent variables (i.e. Bias, Guess rate, and Precision). We used an alpha level of .05. As an exploratory analysis, we also fitted a model including Swap Error, a parameter which accounts for response bias for non-target items. We analyzed the data from this model in the same way as the above-mentioned analyses.

## Results

As depicted in *Figure 2b*, we found an effect of Memory Load on Bias (*F*(3, 174) = 8.94, *p* < .001), such that the bias towards the categorical prototype became stronger with increasing memory load. Specifically, bias was lowest at memory load 1, as compared to higher memory loads (*p* < .001 in all cases), suggesting that VWM predominantly relied on continuous representations at the lowest memory load, before it resorted to categorical representations as memory load increased. Visual inspection of Figure 2b suggests that this pattern was different for younger and older adults. For younger adults, this bias was strongest for intermediate memory loads; in contrast, for older adults, bias increased further in the higher memory load conditions. However, we did *not* find evidence for a significant effect of Age on bias (*F*(1, 58) = 1.01, *p* = .32) or an Age × Memory Load interaction (*F*(3, 174) = 1.14, *p* = .34).

**Figure 2.**
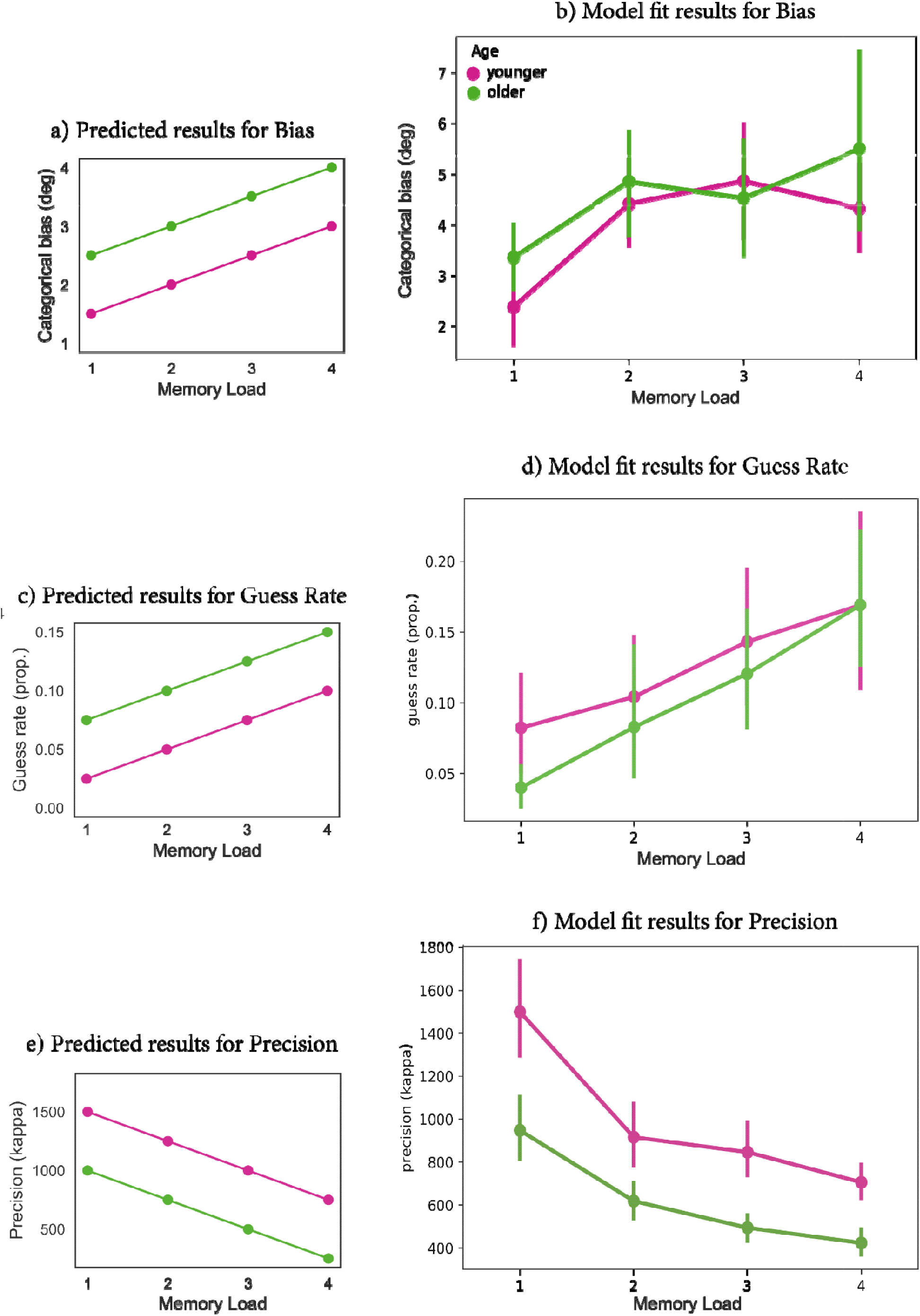
Predicted (on the left) and model fit (on the right) results for Bias, Guess Rate and Precision for both younger and older group.

For Guess Rate, we found an effect of Memory Load (*F*(3, 174) = 33.03, *p* < .001), such that guess rate increased with increasing memory load (*Figure 2d*). We did not find evidence for an effect of Age on guess rate (*F*(1, 58) = 0.56, *p* = .46) or an Age × Memory Load interaction (*F*(3, 174) = 1.13, *p* = .34). For Precision, we found an effect of Memory Load (*F*(3, 174) = 55.02, *p* < .001), such that precision decreased with increasing memory load. Moreover, we found an effect of Age on Precision (*F*(1, 58) = 34.79, *p* < .001), such that older adults were less precise as compared to younger adults (*Figure 2f*). We did not find statistical evidence for an Age × Memory Load on Precision (*F*(3, 174) = 2.51, *p* = .06).

Figure 3 illustrates the response frequency of each memory color for each age group and memory load condition separately. For the younger group, the responses became more centralized towards the category prototypes (represented by the vertical solid lines), and consistent with previous findings (Zhou et al., 2022), this centralization reached a peak at memory load 3. For the older group, the responses were centralized toward the prototypes at lower memory loads; interestingly, their responses became more uniform (and less centralized toward the prototypes) as memory load increased to three. Moreover, Figure 4 shows the distribution of response errors from the memory color for each age group and memory load separately. As reflected in the width of distribution, the overall precision declined with memory load. In particular, this figure shows a strong decline in response precision for older adults at higher memory loads.

**Figure 3.**
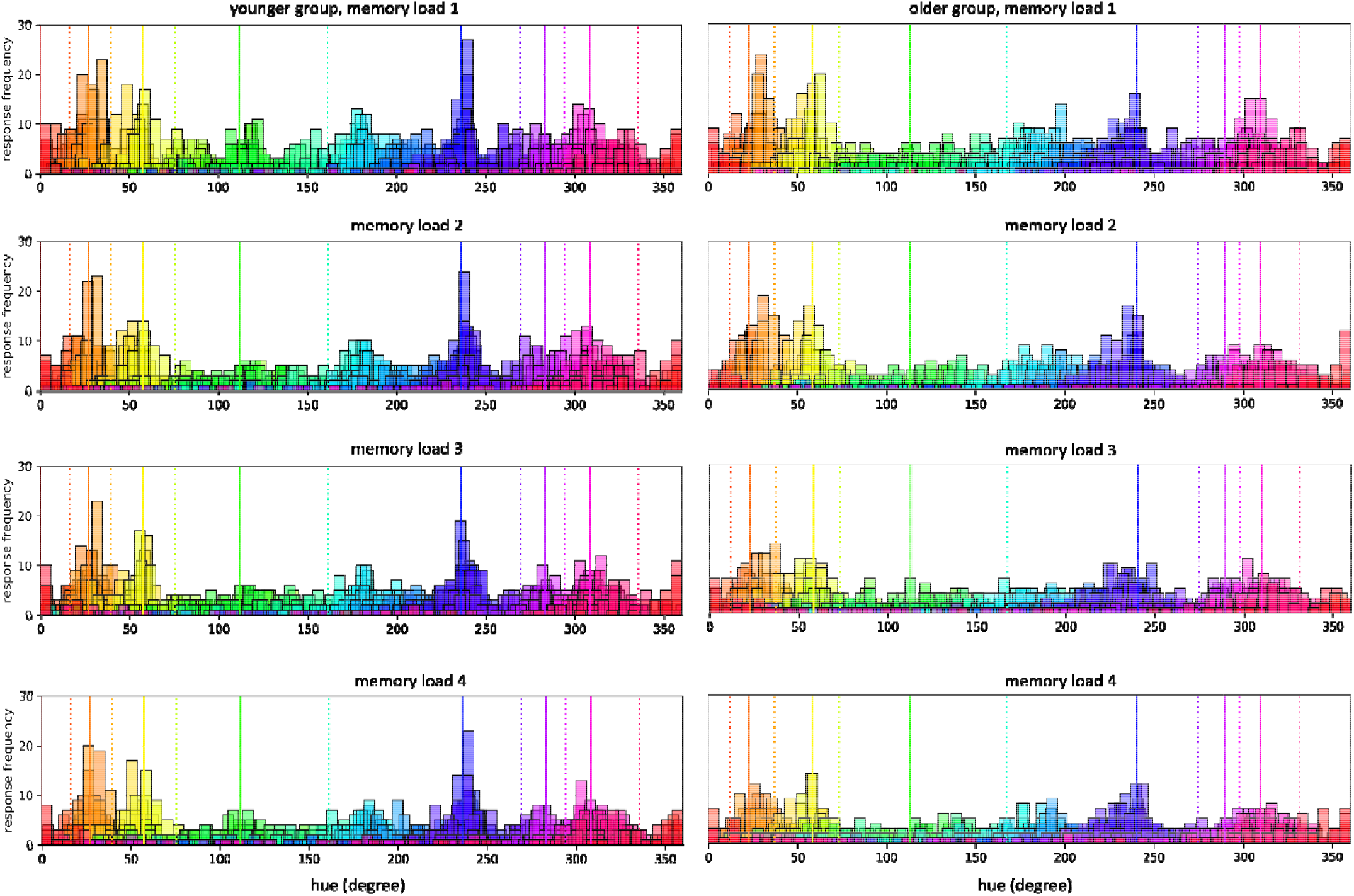
Response frequencies for each hue for the younger and older group. Vertical dotted lines indicate the mean color-category boundaries, and vertical solid lines indicate the mean category prototypes for each age group.

**Figure 4.**
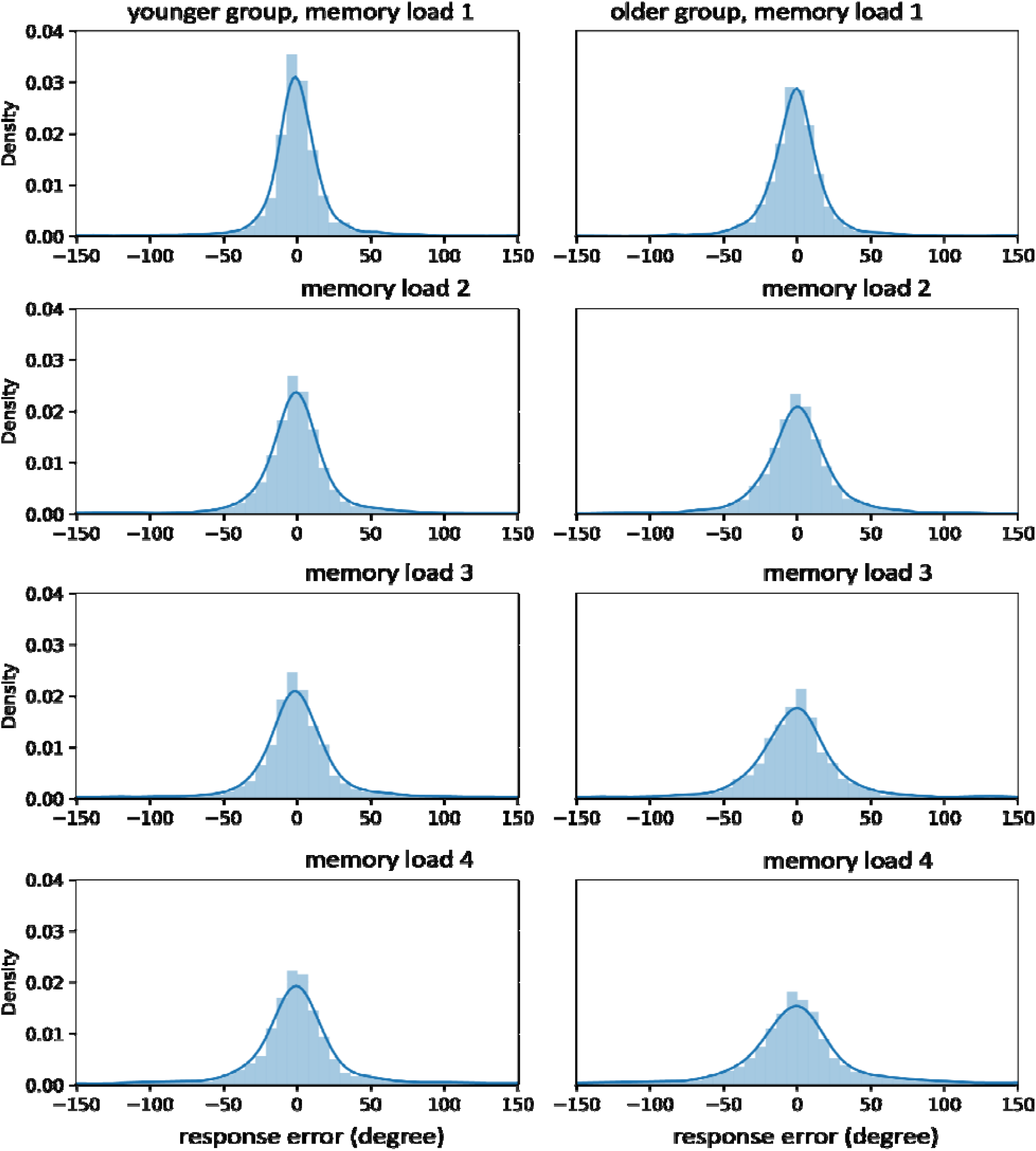
Distribution of response errors for the younger and older group as a function of memory load.

In sum, we found an effect of memory load on bias, such that categorical bias increased with increasing memory load. We did not observe an age effect on bias, suggesting that the extent to which VWM relied on categorical representations was unaffected by age. However, our results suggest that older people likely failed to maintain the colors in VWM, which might contribute to the high biases at higher memory loads.

### Exploratory analysis

In addition, we fitted a model which includes Swap Rate, a parameter that accounts for response bias towards non-target colors (i.e. swap errors). In this model, we found an effect of Memory Load on swap errors (*F*(2, 116) = 8.51, *p* < .001), such that swap errors increased with memory load. There was no effect of Age (*F*(1, 58) = 0.41, *p* = .52) or an Age × Memory Load interaction (*F*(2, 116) = 0.53, *p* = .60) on swap errors. This suggests that swap errors contributed to the deviation of responses as memory load increased; however, it did not contribute to the deviation of responses between age groups.

## Discussion

The present study investigated whether VWM becomes more reliant on categorical representations with age. To this end, we directly measured the extent to which VWM was biased towards category prototypes using an adapted mixture model (the Bias Memory Model; (Zhou et al., 2022). In a delayed estimation task, participants memorized one to four colors (memory load one to four); after a delay, they reported one of the memory colors on a color wheel. We calculated response bias, which reflects the response error in the direction of the category prototype to which the target color belongs; then, we fitted three parameters using the Biased Memory Model: Bias (or categorical bias), Guess Rate, and Precision, to the distribution of response bias.

We found that for both younger and older adults, VWM was biased more strongly towards the category prototype with increasing memory load. Specifically, we found that for both age groups, categorical bias was lowest at memory load 1, as compared to higher memory loads. This suggests that VWM predominantly relied on continuous representations at the lowest memory load, and became predominantly reliant on categorical representations as memory load increased. Interestingly, for younger adults, categorical bias peaked at intermediate-to-high memory loads, whereas for older adults, bias peaked at the highest load. This might reflect different patterns in the way visual information is stored at the higher memory load conditions.

Typically, visual information can be present in two distinct states: within and outside the focus of attention (Cowan, 1998; Oberauer, 2002). When information is presented within the focus of attention, information is represented with sensory details and high precision in VWM (i.e. continuous representations in the current study; (Foster et al., 2017; Harrison & Tong, 2009; Pasternak & Greenlee, 2005). In contrast, when information is presented outside the focus of attention, this information is represented less detailed in VWM (i.e. categorical representations; Bae & Luck, 2019).

In the current study, attentional resources had to be divided over more items as memory load increases. However, since the attentional system has limited capacity, especially in older adults, capacity limitations in attention might have led to more severe consequences on task performance in older adults in the higher memory load conditions (Glisky, 2007; Nagamatsu, Carolan, Liu-Ambrose, & Handy, 2011; Zanto & Gazzaley, 2014). In line with this, evidence has shown substantial age differences in the capacity to divide attention across spatial locations (Greenwood & Parasuraman, 2004; Hüttermann, Bock, Memmert, & Sport, 2012; Lawrence, Edwards, & Goodhew, 2018; Pesce, Guidetti, Baldari, Tessitore, & Capranica, 2005). For example, Lawrence and colleagues (2018) showed that older adults focused their attention more narrowly than younger adults, who were able to distribute their attention over a broader spatial area. Therefore, in the current study, younger adults could still have been able to process visual information at higher memory loads, although this information was stored less precisely, in a more categorical way (see *Figure 3 and 4*). In contrast, older adults might have failed to encode, at least part of visual information with increasing memory load, resulting in lower levels of precision, as reflected in the increasingly uniform response distribution of memory colors (*Figure 3*), and declined memory fidelity *(Figure 4)* from memory load 3.

Although the results suggests that selective spatial attention might be hampered in older adults, evidence supporting age-related changes in the spatial spread of attention is mixed (e.g., Langley, Gayzur, Saville, Morlock, & Bagne, 2011; Quigley, Andersen, & Müller, 2012). Instead, the less precise memory responses in older adults might be related to increased susceptibility to the distractions of non-target information (Cowan, Johnson, & Saults, 2005;

Oberauer, Awh, & Sutterer, 2017). However, our analysis on swap errors of non-target colors showed no difference between the two age groups, therefore this explanation seems less likely.

Alternatively, the observed age-related decline in precision might not be related to the initial encoding of visual information, but to the declined ability in retrieving memory items (Rhodes et al., 2020). In particular, for older adults, once the encoded memory colors in VWM were not highly accessible anymore after a two-second delay period, the precision of VWM items might have decayed substantially. Therefore, when older adults retrieved this information during recall, their responses became less precise as compared to younger adults.

Importantly, despite the age difference in memory fidelity, we found that the extent to which VWM relies on categorical representations was insensitive to age. This is inconsistent with studies using precision or probability as measures of categorical representations in VWM (Forsberg et al., 2020; Pilz et al., 2020). These results, although appear to be counterintuitive and consistent with the ‘dull hypothesis’ (Perfect & Maylor, 2000), might suggest different mechanisms that underlie VWM representations in younger and older adults.

Although we encouraged visualized, non-verbal WM of colors in the current study (see *Stimuli, Design and Procedure*), older participants might have relied on verbal approaches such as labeling (Souza & Skora, 2017) during the VWM task. Using similar methods as in the current study, Forsberg and colleagues (2020) showed that older adults were more likely to spontaneously use verbal labels (such as ‘blue’) to maintain categorical representations. Specifically, for older adults, the estimated capacity of categorical representations in a silent VWM task did not differ from the same task with instructed labeling; however, when labeling was suppressed, this capacity decreased substantially. In comparison, for younger adults, the capacity for categorical representations increased under labeling, whereas it did not change when labeling was suppressed, as compared to the silent task. This suggests that older adults might have been more reliant on verbal WM to compensate for the decreased capacity in visual WM, whereas younger adults were predominantly reliant on visual WM (Aziz et al., 2021; Park & Reuter-Lorenz, 2009; Reuter-Lorenz & Park, 2014). In support of this assumption, neurological evidence has shown that even in visual-only WM tasks, older adults showed considerate bilateral activities in both left and right frontal areas for verbal and visual WM, respectively, whereas younger adults showed only unilateral activities in these areas (Reuter-Lorenz et al., 2000), suggesting that older adults may engage *both* verbal and visual WM to maintain visual information. Therefore, although it appears that older adults showed no difference in the pattern of using VWM representations as compared to younger adults, they likely relied more on verbal WM to compensate for their visual WM capacity. A future direction will be to test the current findings using methods (e.g., EEG and fMRI) that show the recruitment of verbal and visual WM in a straightforward way.

Taken together, the current study replicated the previous findings on the effect of memory load on VWM representations, such that VWM became more reliant on categorical representations with increasing memory load (Bae, Olkkonen, Allred, & Flombaum, 2015; Hardman et al., 2017; Panichello et al., 2019; Pratte MS et al., 2017; Zhou et al., 2022), and extended these findings to older adults. Crucially, our results expanded our understanding of the way visual information is maintained in VWM with aging, highlighting that while memory fidelity was compromised as age increased, the extent to which categorical representations were engaged in VWM maintenance remained unaffected by age.

### Open practices statement

All experimental data, analyses and other materials can be found on the OSF (Open Science Framework): https://osf.io/2u9zh/. A pre-registration of the experiment is available at: https://osf.io/hyxa3.

## Footnotes

1 This criterion deviates from the pre-registered criterion of 2.5 SD. After each participant completed the experiment, we checked the categorization results; if the results seemed to indicate that the participant responded randomly (e.g., the participant chose a greenish color as the prototypical color of blue), we checked whether the participant understood the instructions correctly; if that was not the case, we asked the participant to redo the categorization task. Therefore, we used a stricter criterion than the pre-registered criterion that was chosen based on a previous study (Zhou et al., 2022), in which participants could not redo the categorization task.

